# Whey Protein Phospholipid Concentrate Supplementation Prevents High-Fat Diet Induced Cognitive Impairment in Wistar Rats by Promoting Brain Neuronal Connectivity and Sphingolipid Clearance

**DOI:** 10.1101/2025.04.29.647813

**Authors:** Duncan A. Sylvestre, Nuanyi Liang, James Guinto Galan, Amber Safar, Felipe de Costa Souza, Mitchell C. Bancks, Vedanth Sundaram, Brian W. Scott, Kasey Schalich, Michael Goodson, Jennifer M Rutkowsky, Kristopher Galang, Gulustan Ozturk, David A. Mills, Daniela Barile, Ameer Y. Taha

## Abstract

Whey protein phospholipid concentrate (WPPC), a byproduct of whey protein processing, is high in phospholipids and glycoconjugates which serve as substrates for fatty acids and sugar monomers (e.g. sialic acid) critical to neuronal myelin synthesis in the brain. This led us to hypothesize that WPPC will improve cognitive impairment induced by a high fat (HF) diet by promoting myelin turnover and improving myelin-dependent processes associated with encoding and storing memory. Male Wistar rats were randomized to one of four diets starting at weaning to ∼6.5 months on age: a low-fat (LF) diet containing 10% fat by weight, a HF diet containing 45% fat by weight to induce cognitive impairment, and a HF diet containing either 1.6% or 10% WPPC by weight (n=12 per diet). Rats were subjected to cognitive testing after 2 and 4 months of dietary intervention and then implanted with chronic bipolar electrodes to measure axonal evoked responses within the entorhinal cortex-hippocampal circuitry. Phospholipid and sphingolipid components of myelin were quantified in the hippocampus. There were no significant differences in cognition measured by novel object recognition after 2 months of supplementation. At 4 months, rats on the HF diet performed significantly worse than rats on the LF, HF1.6 and HF10 diets. The beneficial effects of WPPC on cognition were due to a partial reversal in evoked response impairments in hippocampal memory storage. Additionally, hippocampus sphingolipids were higher in rats on the HF diet compared to the LF, HF1.6 and HF10 groups. These findings demonstrate that WPPC prevented cognitive impairment induced by a HF diet by regulating entorhinal cortex-hippocampal circuitries associated with memory storage, though modulating myelin turnover.

## Introduction

Whey protein phospholipid concentrate (WPPC)^1^ is an underutilized phospholipid-rich byproduct derived from the creation of whey protein supplements. It is mostly used as a low value product in animal feed, although some companies are starting to include it in infant formula. WPPC is enriched in oligosaccharides (21% by weight) and phospholipids, which account for approximately 20% of the total lipids^2^. Phospholipids within WPPC are primarily composed of phosphatidylcholine (PC; 36% of total phospholipids) and phosphatidylethanolamine (PE; 38%), although other fractions such as phosphatidylserine (PS; 14%) and phosphatidylinositol (PI; 8%) are present as well.^2^

Phospholipids derived from dairy have been shown to improve cognition in aged rodents^3^. For instance, rats receiving a treatment of freeze dried buttermilk concentrate, containing a lipid composition similar to WPPC (phospholipids made up 14% of total lipids), performed significantly better in the Morris Water Maze test than rats receiving a control diet^3^. In a transgenic model of Alzheimer’s Disease, WPPC at a dose of 3.4g/kg per day showed both improvement at the cognitive level measured by performance in a Morris Water Maze and a Y-maze test, and also an amelioration of beta-amyloid and tau pathology^4^. Similarly, a daily supplement of 15g/d of WPPC in humans with mild cognitive impairment (MCI) for 12 months showed a modest improvement in global cognition^5^. While WPPC or similar milk fractions may slow the progression of cognitive impairment to a limited extent, it is not known whether WPPC prophylactically prevents age-related cognitive impairment.

One reason why WPPC may prevent cognitive impairment is because it is a source of phospholipids and conjugated glycans (i.e. glycoconjugates), which serve as substrates for fatty acids and sugar monomers (e.g. sialic acid) critical for brain myelin synthesis and turnover^6–8^. Myelin synthesis and turnover declines with age, resulting in reduced synaptic connectivity and progressive cognitive decline^9^. Neuronal myelin is primarily composed of 40% phospholipids and sphingolipids containing sugar oligomers or monomers such as sialic acid esterified to a sphingosine backbone^10^. Neither phospholipids nor conjugated glycans cross the blood brain barrier, but they can be acted upon by lipases and glycosidases, respectively, in peripheral tissues or the gut to release free fatty acids and sugar monomers which can enter the brain and assemble there to form neuronal phospholipids and sphingolipids^11,12^.

In the present study, we tested the hypothesis that WPPC prevents cognitive impairment induced by a high-fat diet by promoting connectivity between brain areas involved in learning and memory through myelin lipid remodeling^13,14^. Feeding the HF diet chronically models accelerated aging associated with metabolic syndrome, and results in MCI-like phenotypes over time^15^. Male Wistar rats were randomized from weaning until ∼ 6 months of age to one of four diets: a low fat (LF) control diet containing 10% fat by weight, a high fat (HF) diet containing 45% fat by weight, and a HF diet supplemented with either 1.6% or 10% WPPC by weight. Wistar rats were chosen due to their more rapid development of metabolic syndrome symptoms relative to other strains such as Sprague-Dawley rats^16^. Males were chosen because female rats are more resilient to developing cognitive impairment when placed on a HF diet^17–19^. Cognition was measured by a novel object recognition (NOR) test as a measurement of working memory without primary reinforcement after 2 and 4 months on the diet. Shortly thereafter, the rats were implanted with permanent electrodes in the entorhinal cortex and hippocampus to quantify the evoked response, which reflects the neural activity involved in encoding (entorhinal cortex), transferring and consolidating memory in the hippocampus. This approach measures synaptic connectivity between the two regions and helps to elucidate the flow of information critical for memory formation and storage, as the entorhinal cortex is a nodal point connecting the hippocampus to a variety of other sensory cortices^20–22^, while the hippocampus is a structure critical in memory formation^23^. Following euthanasia, phospholipids and sphingolipids were measured in the hippocampus as markers of neuronal myelin integrity.

## Materials and Methods

### Subjects

Twelve lactating Wistar dams (295-400g), each with a litter of four male pups were purchased from Charles River (St. Constant PQ, Canada), approved for this experimental protocol by the Institutional Animal Care and Use Committee at the University of California, Davis, a USDA-registered and AAALAC International accredited facility. The animals were cared for in accordance with the *Guide for the Care and Use of Laboratory Animals* and the American Association for Laboratory Animal Science Position Statements entitled “Humane Care and Use of Laboratory Animals”, “Alleviating Pain and Distress”, “Scientific Basis for Regulation of Animal Care and Use”, “Determining Laboratory Animal Housing Standards”, and “Standards for Assessing the Quality of Laboratory Rodents.” The animals arrived in two batches staggered by one week. The pups were approximately 14 days old upon arrival. During shipping, two of the litters in the first batch were mixed together in the shipping crate, so the pups for these two litters were randomly separated into the different experimental groups.

The rats were housed in a temperature and humidity-controlled vivarium (23°C, 50% humidity), one litter (6 animals) per cage during lactation. Rats were weaned at 21 days of age and separated into one of four groups – low fat (LF), high-fat (HF), HF +1.6% WPPC by diet weight (HF1.6), or HF +10% WPPC (HF10) by diet weight, at which point the dams were euthanized. Diet composition and ingredients are shown in **Supplementary Table 1**. WPPC was sourced from Milk Specialties Global (Eden Prairie MN, USA). Rats were housed 3 per cage until the animals were 60 days old, at which point they were reduced to 2 per cage, and after exceeding 500g in weight, the animals were singly housed for the remainder of the experiment. Food and water were available *ad libitum,* and food was changed every 2-3 days to limit lipid oxidation in the diets.

Animal weight was measured weekly, and food intake was measured every 2-3 days subtracting the amount of food left in the food hoppers from the amount of food provided to the animals a few days prior.

### Behavioral Testing

To test the effect of the diets on cognition, the animals were subjected to an open field test and the Novel Object Recognition (NOR) test^24^ at approximately 2 and 4 month of age (i.e. 44 days and 126 days respectively since initiation of the dietary intervention at weaning). Animals were brought into a dedicated behavior room with a white noise generator for 30 minutes prior to behavioral testing. The rats were placed in an open field arena (dimensions 70cm x 70cm) for 30 minutes to habituate to the testing environment and record locomotory activity. The next day, this open field habituation was repeated for 30 minutes, after which the animals were introduced to two identical objects (familiarization) for 15 minutes. After an inter-trial interval of at least 60 minutes, the animals were placed back into the arenas for five minutes, but with one of the two identical objects switched for a novel one. All objects were cleaned with ethanol prior to every trial to eliminate any olfactory cues. To eliminate the possibility of recognition of previous objects, two different sets of objects were used when the test was performed at two and four months.

NOR scores were evaluated by measuring how much time each animal’s nose was within a zone slightly larger than the “novel” object relative to the amount of time the animal’s nose was within a similar zone for the “familiar” object, using the Ethovision XT software (Noldus, Leesburg VA). This was used to determine the time spent exploring the novel object relative to the familiar object. The discrimination index was calculated as the ratio of the difference between the time spent interacting with the novel object and the familiar object divided by the sum of the time the animals spent interacting with either object.

### Electrode Implantation

After the NOR tests were completed, bipolar electrodes were implanted into the right entorhinal cortex and hippocampus of each subject. Electrodes were made from two 10mm long stainless steel wires with an 8mm pedestal that had been twisted together and separated slightly at the tip by a scalpel or ruler (Plastics One, cat. # 8IMS3031SPCE, Boerne TX). Implantation was conducted under isoflurane anesthesia (5% for induction, ∼2-3% for maintenance; Fluriso™, VetOne, Boise ID) with stereotaxic techniques according to Gombos et al^25^. Anesthetized animals were shaved on top of the head and the surgical site was disinfected with 7.5% povidone-iodine solution followed by sterile ethanol wipes. Eye ointment (Bausch and Lomb, cat. # AB1336, Stockton CA) was applied to the eyes to prevent them from drying out. Subsequently the incision was made along the interaural line and the periosteum was dissected away. The coordinates for the entorhinal cortex were at 7.5mm posterior to bregma, 4.1mm lateral to midline, and 4.25mm down from the skull, and the hippocampus was 3.5mm posterior to bregma, 2.1mm lateral to midline, and 4.5mm down from the skull^25^. A dental drill was used to drill a hole through the skull at these coordinates. Two jeweler’s screws were placed partially in the skull on the opposite side of the midline to act as an anchor for the implant. Once electrodes were placed, dental cement (Lang Dental Manufacturing, cat. # 1806, Wheeling IL) was used to build the implant and hold everything in place, at which point sutures were applied. Between applications of dental cement, a total of ∼1mL of room temperature sterile 0.9% saline (Hospira, cat. # 0409-4888-03, Lake Forest IL) was applied over the implant site to minimize exothermic heat transfer to surrounding soft tissues. Subsequently, two mL of 0.9% saline was administered subcutaneously immediately after surgery to minimize the risk of dehydration. Each rat received a subcutaneous injection of ketoprofen (Ketofen, 5mg/kg, Zoetis, Union City CA) preoperatively and for two days after surgery for pain management. Rats were transferred to clean cages placed on top of a heat pad after each surgery until the animal was sternal, alert and ambulatory. Animals were given 7-14 days post-implantation to recover, prior to measuring evoked responses as described below. All rats were weighed before and after 3 days and 7 days of surgery, and at the time of euthanasia.

### Evoked Response

To measure the evoked response, each rat was connected to an A-M Systems Model 4100 Isolated High Power Stimulator (A-M Systems, Sequim WA), and an A-M Systems Model 1700 Differential AC Amplifier and Axon Digidata 1550A Low Noise Acquisition System digitizer (Molecular Devices, LLC, San Jose CA). Each rat was placed inside an acrylic box (dimensions: 47cm x 29cm x 30cm). The electrode implant on the animal was connected to the stimulator via a loose wire connector, which allowed the animals to move around freely. Each animal was stimulated at an ascending current of 40µA-1800µA to establish a current response curve, and after a subsequent 30-60 minute interval, an evoked response was taken at the expected 50% current that elicited an evoked response. Stimulation was delivered in a train of 10 biphasic pulses with a duration of 0.1ms each phase, with 10 seconds between pulses. Consequently, each stimulation lasted for approximately two minutes, and the animals were given a minimum of three minutes between trains to rest. The resulting traces consisting of the evoked response, which is a measurement of axonal function, were identified according to the descriptions of Racine and Milgram^26^.

After the last evoked response, the animals were deeply anesthetized with 5% isoflurane and euthanized by transcardial perfusion with room temperature phosphate buffered saline, after collecting blood by cardiac puncture. Tissues (heart, liver, lung, adipose, brain – separated into cerebellum, hippocampus and cortex, spleen, colon contents, ileum) were collected, frozen on dry ice, and stored at -80°C until later analysis. Perirenal, visceral, and interstitial adipose and liver were weighed immediately after dissection.

### Phospholipid and sphingolipid Fatty Acid Measurements

Hippocampal samples were homogenized in 600µL of 0.006% β-hydroxytoluene in methanol. 300µL of this homogenate was then diluted with 700µL of 0.006% β-hydroxytoluene in methanol, and subsequently 2mL of chloroform and 0.75mL of aqueous 1mM disodium EDTA/0.9% KCl was added to reach a final ratio of 4:3:1 chloroform:methanol:aqueous (a “Folch” extraction)^27^. The bottom chloroform layer was transferred to a new tube, and lipids from the original tube were re-extracted using chloroform. The samples were dried and reconstituted in 1mL of 2:1 chloroform/isopropanol. 60µL of this extract was spotted onto silica gel plates (Unisil, silica gel G, 20 x 20cm, Shenzhen China) pre-washed in a glass tank containing 100 mL of 2:1 chloroform/methanol for 1 hour and dried overnight in a vacuum oven at 80℃. Phospholipid classes were separated using a solvent mixture of 60:50:4:1 (v/v) chloroform:methanol:water:glacial acetic acid for approximately 1 hour. Lipid bands were visualized with 0.02% 2’7’-dicholorofluorescein in methanol under UV light, scraped and placed into fresh tubes containing C19:0 PC as an internal standard, and 0.4mL of toluene. Samples were then transesterified according to Zhang et al^28^, with 3mL of methanol and 600µL of 8% HCl in methanol at 90°C for one hour.

### Dietary and serum fatty acid analysis

To analyze dietary fatty acids, one pellet from each diet was ground to a powder and 50 mg of powder was subjected to the same double “Folch” extraction with 3mL of 2:1 chloroform:methanol and 0.75mL of aqueous 1mM disodium EDTA/0.9% KCl at room temperature and subsequent transesterification. Serum total fatty acids were measured in the same manner, using approximately 25µL of serum. We attempted to measure fatty acids in erythrocytes, but after Folch extraction and transesterification, only small inconsistent peaks were seen on the gas-chromatogram due to extensive clotting. Erythrocytes were therefore not further analyzed.

### Gas Chromatography Analysis of Fatty Acid Methyl Esters

Transesterified samples were analyzed on a Perkin Elmer Clarus 500 Gas Chromatograph system (Perkin Elmer, Shelton, CT, USA) under the following parameters: Initial oven temperature of 80℃ for two minutes, then ramping to 185℃ at 10℃ per minute, then ramping to 249℃ at 6℃ per minute, and holding at this temperature for 42 minutes. Data were expressed as absolute concentrations (mg fatty acid per gram of tissue, or mg fatty acid per mL of serum).

### Statistical Analysis

All data are represented as average ± SD. Body weight data between weaning and the beginning of surgeries was evaluated by two-way repeated measures analysis of variance (ANOVA), while food intake during this time was evaluated by mixed-effects ANOVA. Behavioral data and evoked responses were evaluated by one-way ANOVA followed by Tukey’s post-hoc test. Additionally, a multiple linear regression model was used with evoked response as the output, and diet and time since behavior (aliased to total diet exposure) as confounding variables. Residuals from this model were analyzed with a single-factor ANOVA followed by Tukey’s post-hoc test.

## Results

### Dietary Fatty Acids

**Supplementary Table 2** shows the percent fatty acid composition of the LF, HF, HF1.6 and HF10 diets. As shown, the LF diet had a greater proportion of linoleic acid and linolenic acid, than the HF diets with or without WPPC. Conversely, palmitate, stearate and oleic acid were higher in the HF diet with or without WPPC, compared to the LF diet.

### Postoperative Attrition

Fifteen (31%) rats did not have an evoked response measurement mostly due to electrode detachment after surgery (3 LF, 5 HF, 2 HF1.6 and 5 HF10 out of 12 per group), however, tissues were still collected from eight of these animals (1 LF, 3 HF, 1 HF1.6 and HF10). The reason for electrode detachment is likely because we used 2 screws instead of to stabilize the implants. These rats were therefore immediately euthanized. Additionally, three animals (one animal each in the HF, HF1.6, and HF10 groups) did not have an evoked response measurement due to electrical noise during measurement that made establishing a current response curve difficult. One animal from the HF10 diet group was missing fatty acid data from the hippocampus as the sample was lost during extraction due to a tube shattering. One serum sample from the HF diet group was missing serum fatty acid data because the serum had congealed after centrifugation, making it difficult to separate the serum from the erythrocytes.

### Dietary Intake and Body Weights

We observed a significant difference in animal body weight starting at approximately 80 days of age (**Figure 1**), in which all three HF diet groups (with or without WPPC) gained significantly more weight than the LF animals; this weight difference persisted up until the beginning of surgeries, when weekly weight monitoring stopped and the animals were weighed was only measured when the rats underwent surgery.

**Figure 1:**
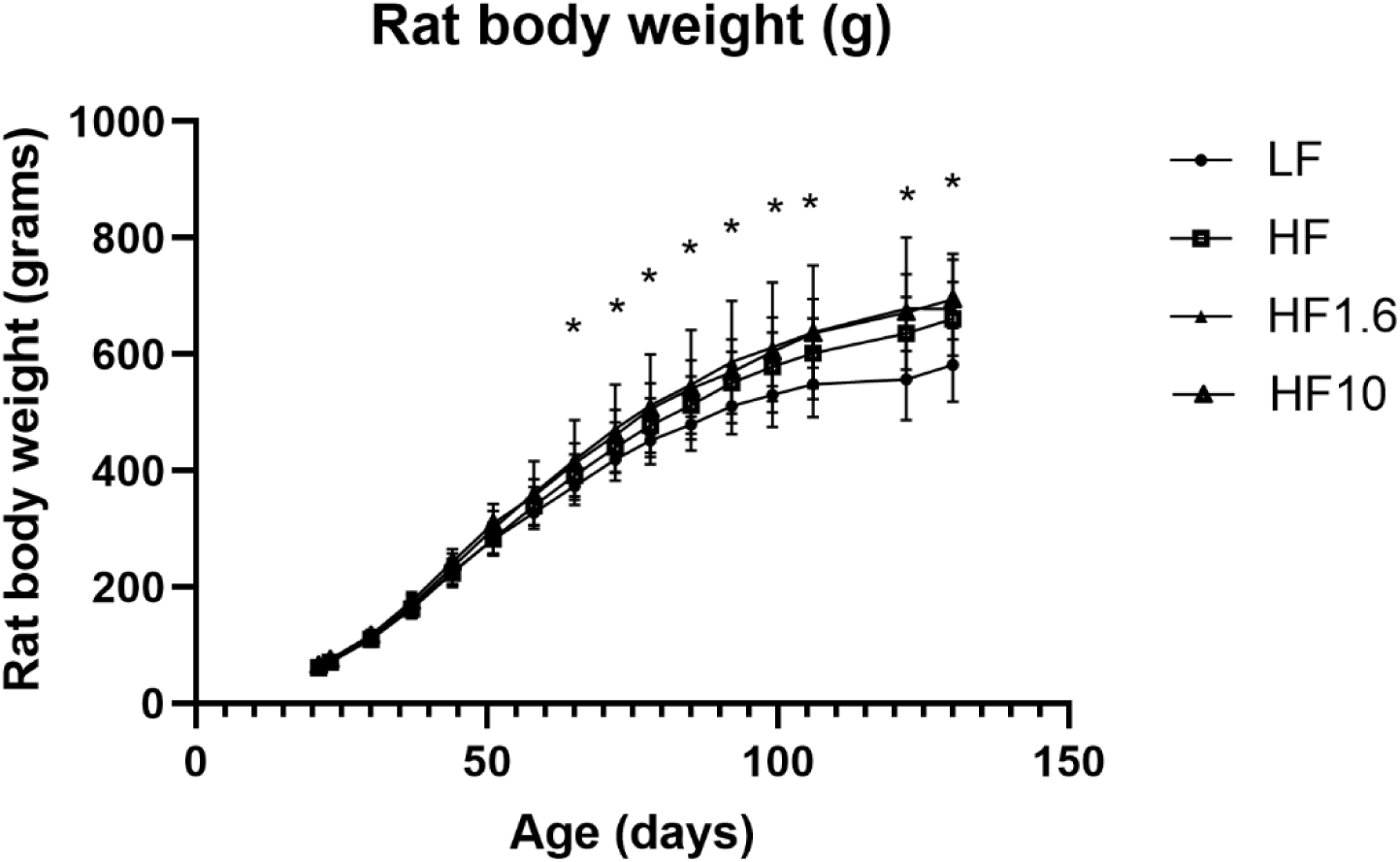
Two-way repeated measured ANOVA of mean body weights of male Wistar rats (n=12 per group) placed on a diet low in fat (10%, LF), high in fat (45%, HF), or high in fat supplemented with 1.6% or 10% whey protein phospholipid concentrate (HF1.6 and HF10 respectively). The LF diet groups were significantly lower than all three groups at each timepoint by Tukey’s post-hoc test, as indicated with the asterisks.

**Figure 2** shows the intake pattern of the rats in each group expressed as both cumulative caloric intake (**Figure 2A**) and as the average caloric intake per day in each 2-3 day feeding period (**Figure 2B**). Starting at approximately 40 days after initiation of dietary intervention the cumulative caloric intake for the three HF groups began to diverge from the intake of the LF group (**Figure 2A**) peaking at a difference of approximately 2000kcal more consumed in the HF groups than the LF group after 110 days of dietary intervention (**Figure 2A**). Absolute food intake (g/day), however, did not differ significantly between the groups over time (**Figure 2B**).

**Figure 2:**
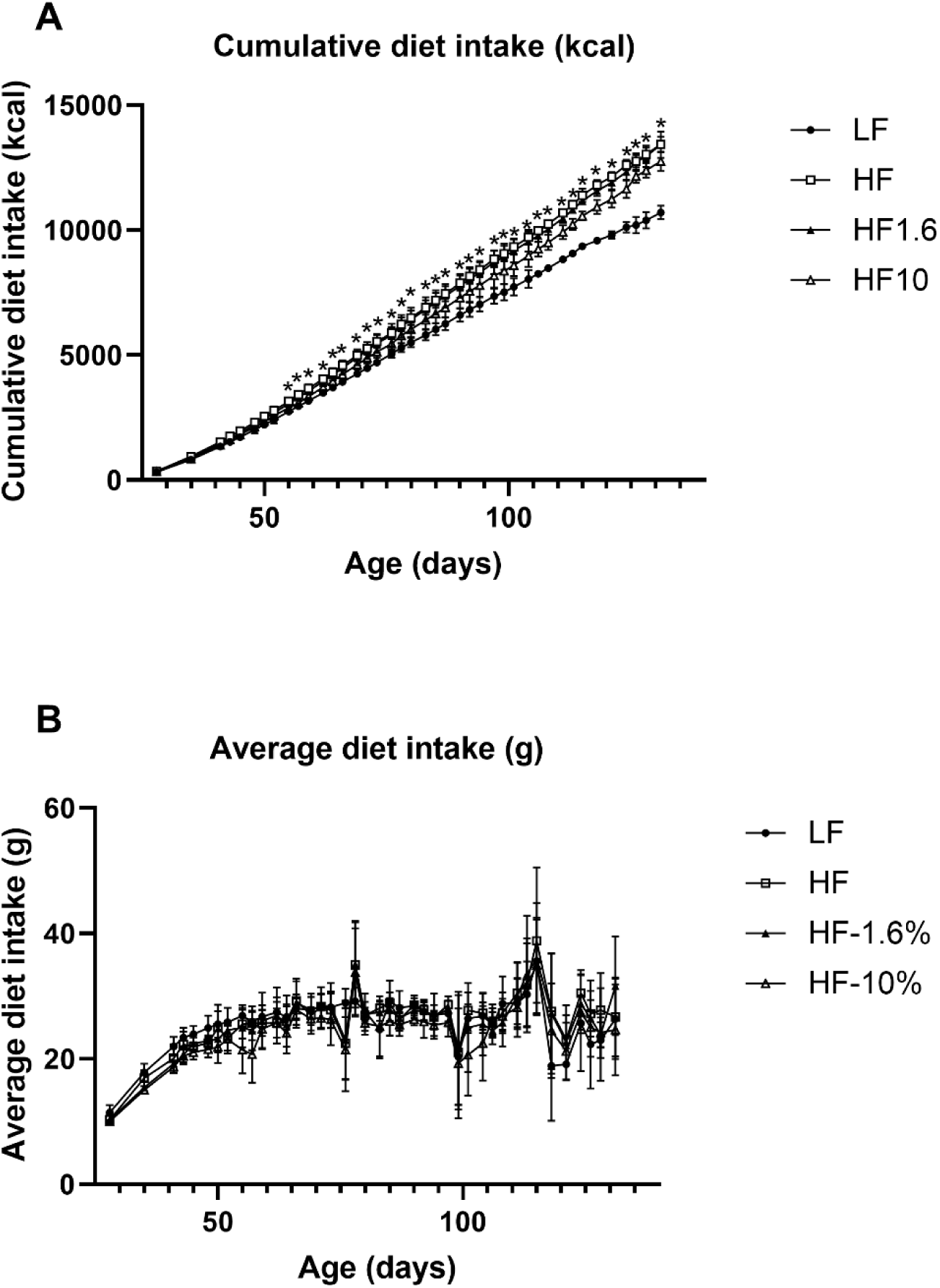
Two-way ANOVA with Mixed Effects Analysis of cumulative kcal consumed (**A**) and average grams of diet consumed (**B**) of male Wistar rats (n=12 per group) placed on a diet low in fat (10%, LF), high in fat (45%, HF), or high in fat supplemented with 1.6% or 10% whey protein phospholipid concentrate (HF1.6 and HF10 respectively). An asterisk above a point indicates a significant difference by Tukey’s post-hoc test at that time point.

### Postoperative Dietary Intake and Weight Loss

Over the span of the seven to fourteen day postoperative period, we observed that animals in the HF group lost approximately twice as much weight relative to the LF group (**Figure 3A**, p=0.0359), despite no significant differences in food intake during this period (**Figure 3B**), with the HF1.6% group weighting significantly more than the LF group at sacrifice (p=0.0455) and the HF10% group trending higher than the LF group (p=0.0869). To explore the cause of these differences, body and tissue (adipose and liver) weights were measured at euthanasia (**Supplementary Figure 1**). As shown, the data confirmed that the HF1.6 group had significantly more body weight compared to the LF group, and this appears to be driven by gains in perirenal and visceral adipose tissue. The HF10 group trended in the same direction, although differences were not statistically significant, likely due to the lower sample size.

**Figure 3:**
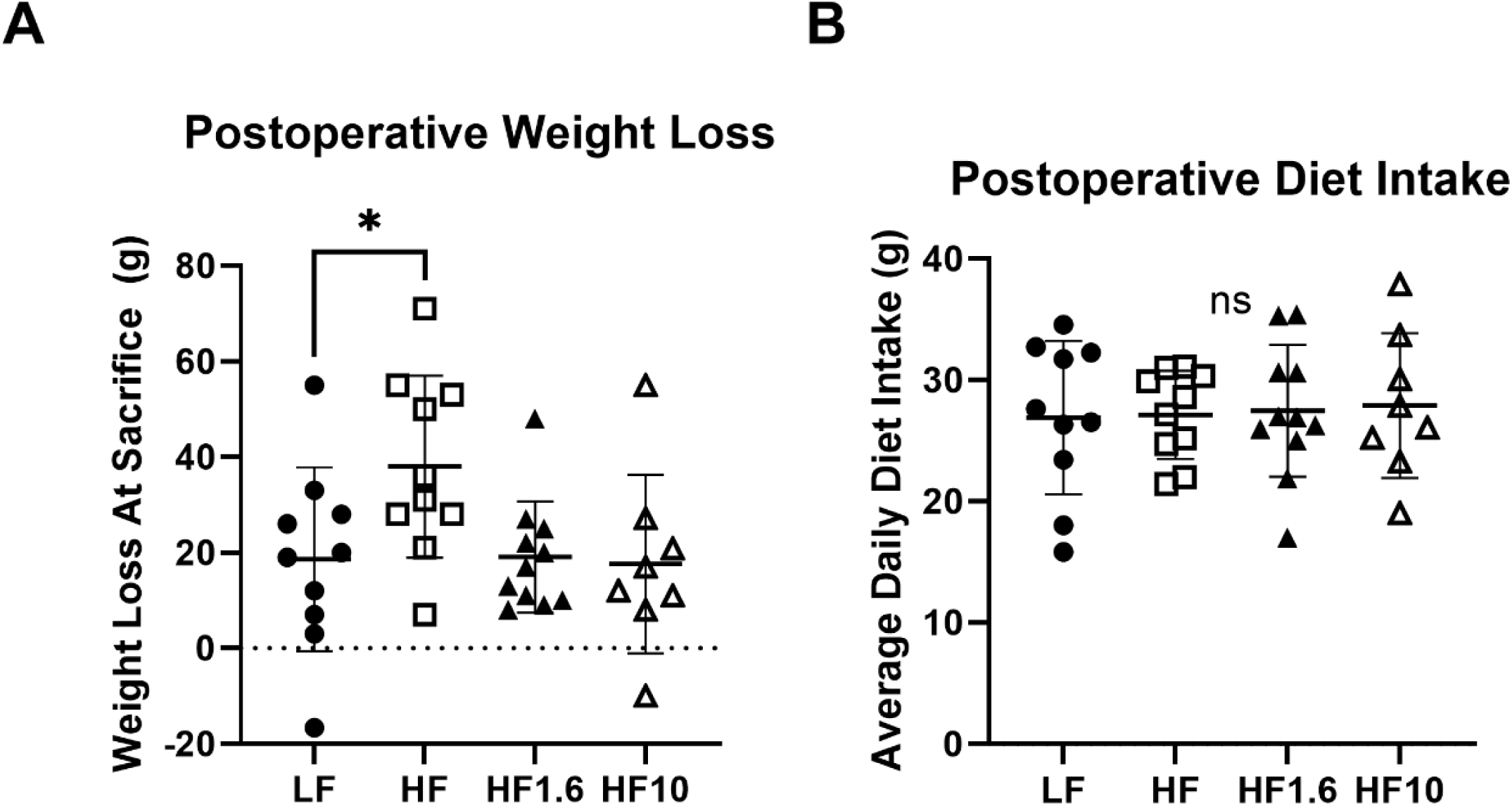
One-way ANOVA of postoperative weight loss calculated by subtracting weight at sacrifice from preoperative body weight (**A**) and average daily dietary intake over the postoperative period (**B**) of male Wistar rats (LF n=9, HF n=7, HF1.6 n=10, HF10 n=6) placed on either a low fat diet (LF, 10% fat by weight), high fat diet (HF, 45% fat by weight), or a high fat diet supplemented with 1.6% or 10% WPPC by weight (HF1.6 and HF10 respectively). One-way ANOVA followed by Tukey’s post-hoc test revealed that the HF group experienced significantly more weight loss during the postoperative period than the other three groups, while there was no significant difference in the amount of food consumed.

### Novel Object Recognition and Open Field Test

There was no significant difference in the discrimination index between the groups when measured at approximately 2 months of dietary intervention (**Figure 4A**). However, after 4 months, we observed that rats fed the HF diet performed significantly worse than the LF, HF1.6%, and HF10% groups (**Figure 4B**; p=0.0014). These results indicate that the HF diets containing 1.6% and 10% WPPC prevented cognitive impairment induced by the HF diet, relative to the LF diet. Due to software data corruption, we were unable to analyze data for the open field at 2 months of dietary intervention. There was no significant difference in the performance on the open field test performed at 4 months (**Figure 4C**), measured by total distance traveled, indicating that there were no differences in mobility between the animals in any group.

**Figure 4:**
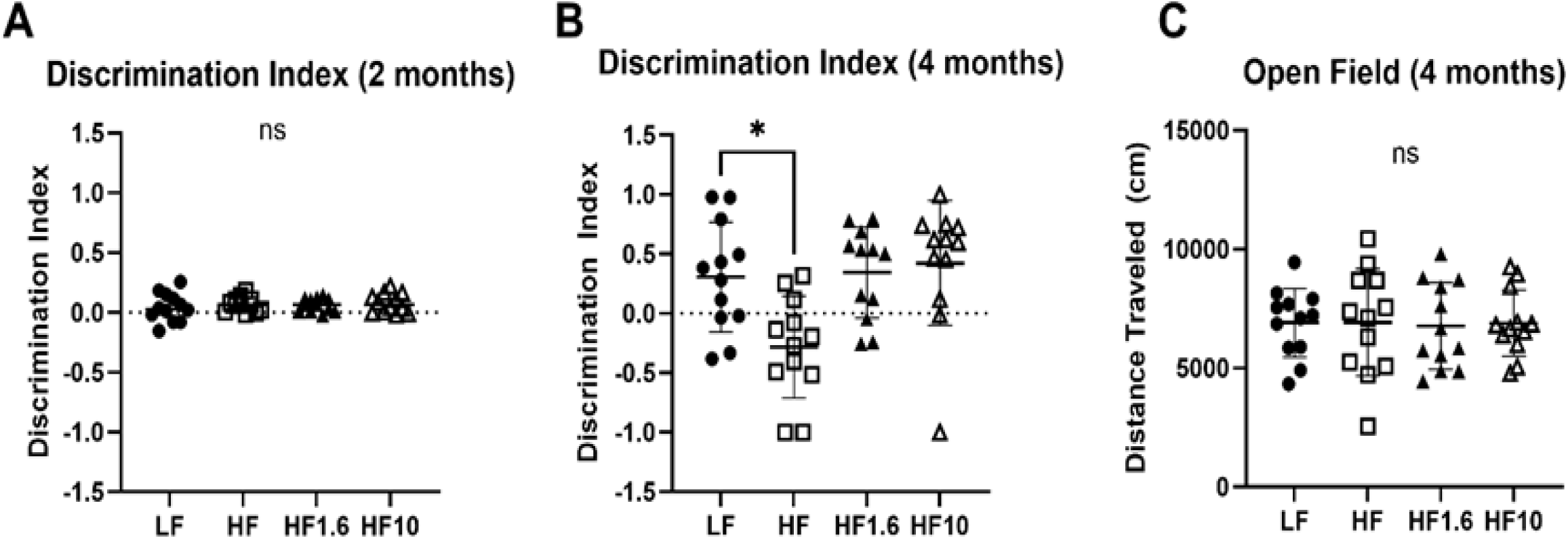
One-way ANOVA of novel Object Test results of male Wistar rats (n=12 per group) placed on a diet low in fat (10%, LF), high in fat (45%, HF), or high in fat supplemented with 1.6% or 10% whey protein phospholipid concentrate (HF1.6 and HF10 respectively) when performed at approximately 2 months (**A**) and 4 months (**B**) of supplementation. One-way ANOVA followed by Tukey’s post-hoc test revealed that animals on the HF diet performed significantly worse at 4 months than LF, HF1.6, and HF10, while there was no difference at 2 months. At 4 months, there was no significant difference in mobility on the Open Field test performed prior to the novel object test on the same cohort (**C**).

### Evoked Response measurements

A representative trace of evoked response following stimulation is shown in **Supplementary Figure 2**. The 50% maximal evoked responses amplitude (“ED50”) at baseline, and ratio of the amplitude following stimulation at the ED50 after 30-60 minutes of obtaining baseline, relative to baseline ED50, did not differ significantly between the groups (**Supplementary Figure 3**).

Because the surgeries were performed over the span of 2 months (i.e. between 4 to 6 months after diet initiation), linear regression was performed to correct for total diet exposure. Multiple regression analysis with main effects only showed a negative relationship between the ratio of the maximal evoked response amplitude (at 30-60 minutes relative to baseline) and the time interval between duration on the diet and stimulation (β = -0.2112±0.1424, p=0.0062). A separate model including a two-way interaction between group and time since behavior showed this negative relationship in the HF (- 0.03384±0.01155, p=0.0073) and to a lesser extent, in the HF1.6% diets (- 0.02624±0.01063, p=0.0211), but not the HF10% diet. Residual plots controlling for diet exposure time for the ED50 and ratio of evoked responses 30 to 60 minutes relative to the ED50 baseline are presented in **Figures 5A and 5B**, respectively. A one-way ANOVA followed by post-hoc Tukey test was performed to highlight differences between the groups. As expected, no significant differences were seen in the ED50 as shown in **Figure 5A** (since the regression showed no significant effects). However, the HF diet appeared to reduce the ratio of the evoked response amplitude compared to the LF diet, though the residuals trended towards statistical significance between the HF and LF groups (P=0.09 by Tukey’s post-hoc test) but not between the LF and HF1.6 and HF 10 groups (P>0.05) (**Figure 5B**). Overall, the data suggest that WPPC partially prevented impairments in entorhinal cortex-hippocampal circuitries associated with memory storage in rats on the HF diet, but not with circuitries associated with encoding memory.

**Figure 5:**
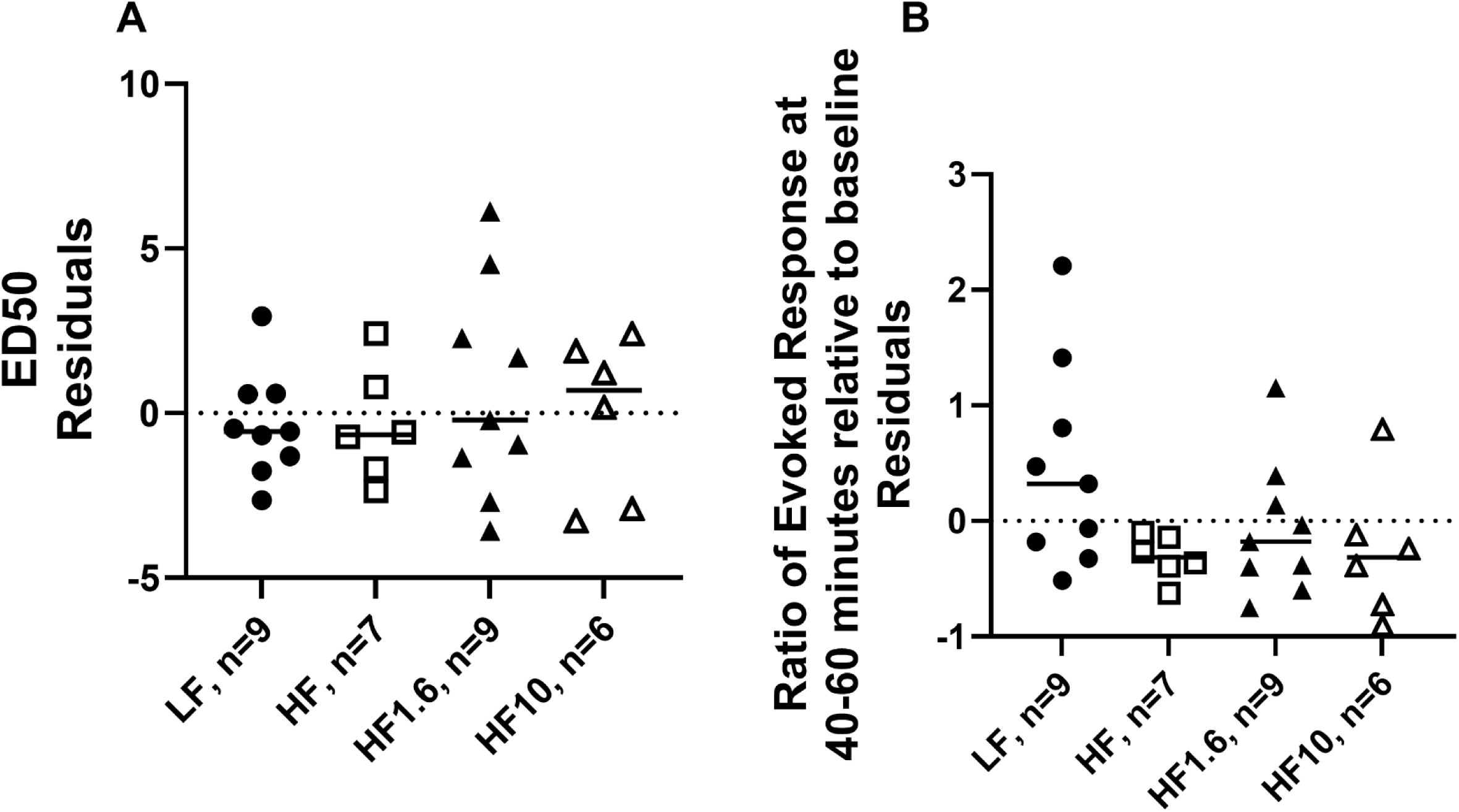
Comparison by one-way ANOVA of the residuals of the 50% evoked response amplitude (ED50) (**A**) and the ratio of evoked response at 30-60 minutes relative to baseline ED50 (**B**), from a multiple regression model controlling for total diet exposure time for rats consuming a LF, HF, HF1.6 or HF10 diets. One-way ANOVA followed by Tukey’s post-hoc test showed no significant difference in residuals across groups, but there was a trend towards significance between the LF and HF groups (p=0.09) for the evoked response ratio (**B**)

### Hippocampal Phospholipids

We observed no significant differences hippocampus phospholipid fractions between the groups (**Supplementary Table 3**). In the sphingolipid fraction (found mostly in myelin), we observed a 140% increase in DHA (p=0.0207) in the HF group relative to the LF diet which was statistically significant (**Figure 6**). These differences were not observed in rats on the HF diet with 1.6% or 10% WPPC. All measured fatty acid concentrations in sphingolipids can be found in **Supplementary Table 3**. As shown, there were similar trends as observed for DHA in other fatty acids, but these changes did not reach statistical significance.

**Figure 6:**
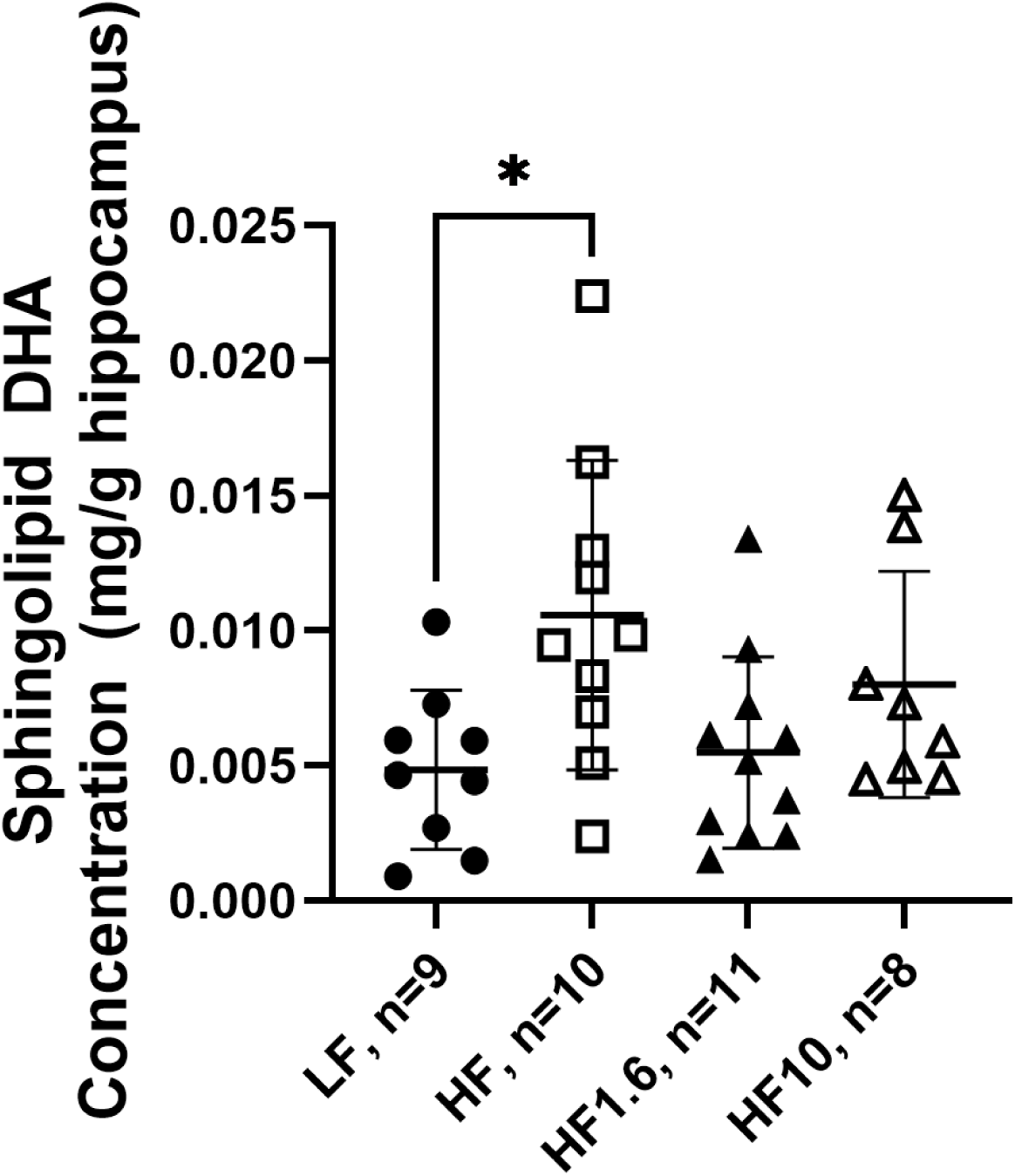
Concentration (mg/g) of DHA in the sphingolipid fraction of the hippocampus of rats consuming a LF, HF, HF1.6 or HF10 diet. The concentration of DHA was significantly higher in the HF group than the LF group (p=0.0207) by one-way ANOVA followed by Tukey’s post-hoc test.

### Serum Fatty Acids

Palmitoleic acid was significantly elevated in the serum of rats on the HF, HF1.6%, and HF10% diets relative to rats on the LF diet (p=0.0002). Additionally, eicosapentaenoic acid was lower in HF, HF1.6, and HF10 animals (p=0.0029) while docosahexaenoic acid trended lower in the HF10 animals (p=0.0804) (**Figure 7**).

**Figure 7:**
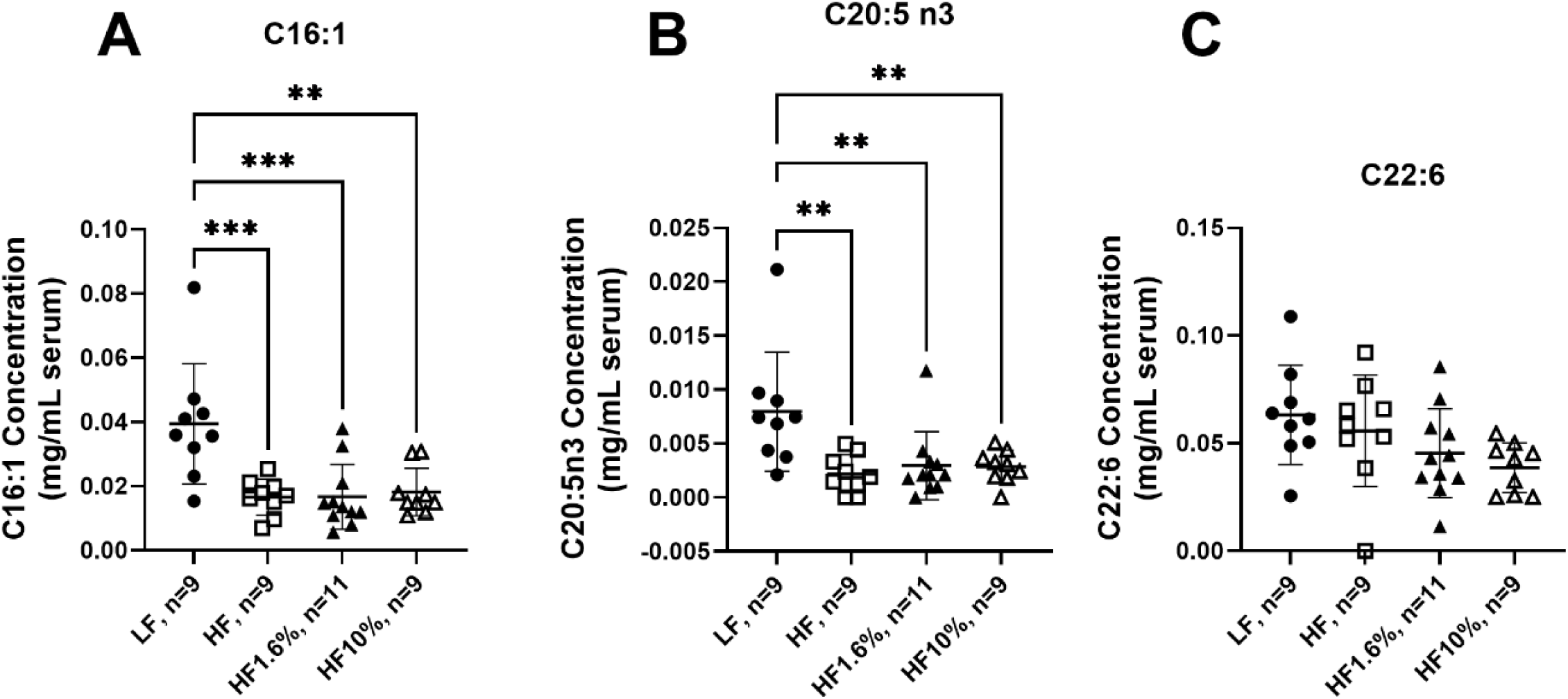
Concentration (mg/mL) of palmitoleic acid (16:1, **A**), EPA (20:5n3, **B**), and DHA (22:6n3, **C**) in the serum of animals on a LF, HF, HF1.6, or HF10 diets. One-way ANOVA followed by Tukey’s post-hoc test showed that16:1 (p= 0.0002) and EPA (p=0.0029) were significantly lower in all HF groups than in the LF group, while DHA (p=0.0804) trended lower in the HF10 group relative to the LF group.

## Discussion

In the present study, supplementing rats with WPPC at 1.6% of 10% (by weight), for 4 to 6 months prevented cognitive impairment induced by a HF diet. WPPC at either doses partially reversed entorhinal cortex-hippocampus evoked response depressions induced by the HF diet and prevented sphingolipid accumulation in the hippocampus. Our findings suggest that the cognitive benefits of WPPC are linked to improvements in neuronal circuities associated with memory storage in the hippocampus, due to enhanced myelin (sphingolipid) turnover.

Rats consuming the HF diet demonstrated significantly worse performance on the NOR test after 4 months, than rats on either a LF diet or HF diet supplemented with either concentration of WPPC (**Figure 4B**). The NOR test is a measurement of working memory involving engagement of the hippocampus for object recency as well as prefrontal cortex and perirhinal cortex for object preference^24,29,30^. Consistent with these observations, we also found that WPPC partially reversed reductions in entorhinal-hippocampus evoked response amplitude ratios caused by the HF diet relative to the LF diet (**Figure 5B**). A reduction in the evoked response ratio due to the HF diet indicates impaired memory storage, as opposed to an impairment in encoding, which would have been represented by a reduction in the baseline evoked response. Thus, our findings suggest that the cognitive benefits of WPPC are likely linked to improvements in entorhinal cortex-hippocampus neuronal circuitries that enable rats to store memory.

Rats consuming the HF diet had significantly more DHA-sphingolipids than rats on the LF diet and rats on the HF diet supplemented with WPPC at 1.6 or 10% (**Figure 6**). The accumulation of sphingolipids in the hippocampus of rats on the HF diet reflects impaired turnover (i.e. synthesis and clearance) of myelin, where sphingolipids are enriched. This finding is in agreement with a prior study which showed that a HF diet promotes sphingolipid accumulation in rat brain^31^. Adding WPPC to the HF diet prevented the accumulation of sphingolipids in our study, suggesting that WPPC promoted sphingolipid clearance to the same levels seen in the LF control group. Notably, sphingolipid accumulation in the brain is seen in lysosomal storage disorders such as acid sphingomyelinase deficiency and Neiman Pick’s disease, where the lack of sphingolipid clearance has been linked to neurodegeneration and cognitive impairment ^32,33^. The HF diet appears to initiate similar phenotypes in our rat model, and this is reversible by WPPC, suggesting that WPPC components target myelin turnover in the brain.

The effects of WPPC on the brain are likely mediated, at least in part, by phospholipids and oligosaccharide glycoconjugates previously characterized in WPPC^2^. Although phospholipids do not cross the blood brain barrier to an appreciable extent, they act as substrates for free fatty acid release in plasma, and supply to the brain (free fatty acids passively enter the brain)^12^. WPPC also contains glycoconjugates which may be released through gut bacteria to provide sialic acid and other sugar monomers^11^ that can be utilized towards myelin synthesis within the brain. Since the effects of WPPC were seen in sphingolipids which are mainly found in myelin, our data suggest that WPPC likely acts by providing key fatty acid and sugar substrates for maintaining myelin turnover. This requires direct validation through tracer experiments in future studies. Future research could also examine alterations in the gut microbiota composition and function in the four study groups.

Animals that consumed the HF diet lost more weight 7-14 days post-operatively compared to rats on the LF diets. These changes in weight loss were prevented in rats on the HF1.6% and HF10% groups. As there were no or small differences in the perirenal and visceral adipose tissue between the three HF groups and the LF group at the time of euthanasia (**Supplementary Figure 3**), this suggests that the lost weight was possible in rats on the HF diet due to sarcopenia. This was prevented by the addition of WPPC into the diet at either dose. The causes of this incidental finding remain unknown. Postoperative weight loss represents a significant health risk as surgeries place the body in a hypermetabolic state^34^. Preventing or mitigating surgery-related weight loss is a field of ongoing research, however, few investigators have evaluated the effects of preoperative nutritional interventions^35^. WPPC represents a potential nutritional intervention that could significantly improve management of postoperative weight loss.

Serum fatty acids were measured to identify fatty acid markers of WPPC intake. We did not observe a marker that differentiates the WPPC dietary groups from the others. However, we saw an increase in palmitoleic acid (16:1n-7) and a reduction in long-chain omega-3 eicosapentaenoic acid and docosahexaenoic acid in rats on the 3 HF diets compared to LF controls. The increase in palmitoleic acid is likely due to the greater dietary composition of palmitic acid originating from lard; dietary palmitate can be elongated to palmitoleic acid in the liver, resulting in greater serum levels as seen in this study. Similarly, the reduction in long-chain omega-3 polyunsaturated fatty acids is likely due to the lower levels of alpha-linolenic acid in the HF diets, which would limit substrate availability for liver elongation-desaturation ^36,37^. It is unlikely that these serum fatty acid changes impacted our brain outcomes, as no significant changes were observed between rats on the HF diet compared rats on the HF diet containing WPPC.

A limitation of the present study is that the diets were not isocaloric, which means that the differences between the LF and HF groups may be due to excess energy intake, rather than the fat content of the diet specifically, though the three HF diets were isocaloric relative to each other. Additionally, the behavioral measurement of cognition that we used, the NOR test, is a measurement of working memory while the entorhinal-dentate pathway that we stimulated to acquire an evoked response is a measurement of spatial memory^38^. While this demonstrates effects in multiple forms of cognition, we did not perform a behavioral measurement of spatial memory such as a Barnes Maze, though we did approximate a measurement of pure recall by comparing the ratios of baseline evoked responses to their values 30-60 minutes afterwards. Additionally, due to electrode displacement after surgery some animals only have a NOR test result and not an evoked response, which could represent a source of survivorship bias.

### Conclusion

Supplementation of WPPC prevented HF-diet induced cognitive impairment to working memory, by promoting lipid clearance within myelin and neuronal connectivity between the entorhinal cortex and hippocampus, leading to improved memory consolidation. While some of the biological actions and mechanisms of WPPC have been elucidated in this study, future experiments should investigate whether the observed effects are mediated through liver or gut release of fatty acids and sugar monomers derived from phospholipids and glycoconjugates found in WPPC.

## Abbreviations List

WPPC: Whey Protein Phospholipid Concentrate
PC: Phosphatidylcholine
PE: Phosphatidylethanolamine
LF: Low Fat
HF: High Fat
HF1.6%: High Fat + 1.6% Whey Protein Phospholipid Concentrate
HF10%: High Fat + 10% Whey Protein Phospholipid Concentrate
NOR: Novel Object Recognition

## Acknowledgements

We would like to thank Dr. Lindsey Ormond and Milk Specialties Global (Eden Prairie, MN, USA) for providing the WPPC used in this study. This study was funded by Dairy Management Inc. grant number 3105-1.

## Author Credit statement

Duncan Sylvestre: Investigation, Formal Analysis, Writing – Original Draft;

Nuanyi Liang: Methodology, Investigation, Formal Analysis;

James Galan: Formal Analysis;

Amber Safar: Formal Analysis;

Felipe de Costa Souza: Investigation;

Mitchell Bancks: Investigation;

Vedanth Sundaram: Validation;

Brian Scott: Formal Analysis; Conceptualization;

Kasey Schalich: Validation;

Michael Goodson: Investigation;

Jennifer Rutkowsky: Resources, Methodology;

Kristopher Galang: Investigation;

Gulustan Ozturk: Investigation;

David Mills: Supervision;

Daniela Barile: Funding acquisition, Conceptualization, Resources, Supervision;

Ameer Taha: Project administration, Conceptualization, Supervision, Writing – Review and Editing, Resources

## Notes

### Competing Interest Statement

The authors have declared no competing interest.

