## Supplementary Material for "Whey Protein Phospholipid Concentrate Supplementation Prevents High-Fat Diet Induced Cognitive Impairment in Wistar Rats by Promoting Brain Neuronal Connectivity and Sphingolipid Clearance"

**Supplementary Table 1**: Ingredients for Low Fat (LF), High Fat (HF), High Fat +1.6% WPPC (HF1.6) and High Fat +10% WPPC (HF10) diets.

|  | ***LF WPI Cont. Diet*** | |  | ***45% HF WPI Cont Diet*** | |  | ***45% HF + 1.6% WPLC Diet*** | |  | ***45% HF + 10% WPLC Diet*** | |
| --- | --- | --- | --- | --- | --- | --- | --- | --- | --- | --- | --- |
| *%* | *gm* | *kcal* |  | *gm* | *kcal* |  | *gm* | *kcal* |  | *gm* | *kcal* |
| Protein | 19.9 | 20.0 |  | 24.7 | 20.0 |  | 24.7 | 20.0 |  | 24.5 | 20.0 |
| Carbohydrate | 72.8 | 70.0 |  | 47.0 | 35.0 |  | 47.0 | 35.0 |  | 46.7 | 35.0 |
| Fat | 4.4 | 10.0 |  | 24.7 | 45.0 |  | 24.3 | 45.0 |  | 22.0 | 45.0 |
| Total |  | 100.0 |  |  | 100.0 |  |  | 100.0 |  |  | 100.0 |
| kcal/gm | 3.98 |  |  | 4.94 |  |  | 4.93 |  |  | 4.91 |  |
| *Ingredient* | *gm* | *kcal* |  | *gm* | *kcal* |  | *gm* | *kcal* |  | *gm* | *kcal* |
| Isopure WPI | 210 | 838 |  | 210 | 838 |  | 201 | 803 |  | 153 | 613 |
| Corn Starch | 533 | 2131 |  | 61 | 245 |  | 61 | 245 |  | 61 | 245 |
| Maltodextrin 10 | 125 | 500 |  | 125 | 500 |  | 125 | 500 |  | 125 | 500 |
| Sucrose | 38 | 223 |  | 143 | 642 |  | 143 | 642 |  | 143 | 642 |
| Lactose | 2.41 | 9.64 |  | 2.41 | 9.64 |  | 2.03 | 8.11 |  | 0.00 | 0.00 |
| Cellulose | 50 | 0 |  | 50 | 0 |  | 50 | 0 |  | 50 | 0 |
| 101952 Dried Whey Cream NK | 0.0 | 0.0 |  | 0.0 | 0.0 |  | 13.6 | 68.0 |  | 85.7 | 428.6 |
| Soybean Oil | 25 | 225 |  | 25 | 225 |  | 25 | 225 |  | 25 | 225 |
| Lard | 22 | 194 |  | 185 | 1661 |  | 181 | 1631 |  | 163 | 1468 |
| AIN-93G Mineral Mix | 35 | 30.9 |  | 35 | 30.9 |  | 35 | 30.9 |  | 35 | 30.9 |
| AIN-93 Vitamin Mix | 10 | 39.2 |  | 10 | 39.2 |  | 10 | 39.2 |  | 10 | 39.2 |
| KH_2_PO_4_ | 0.33 | 0 |  | 0.27 | 0 |  | 0.27 | 0 |  | 0.27 | 0 |
| Choline Bitartrate | 2.5 | 0 |  | 2.5 | 0 |  | 2.5 | 0 |  | 2.5 | 0 |
| TBHQ, antioxidant | 0.014 | 0 |  | 0.014 | 0 |  | 0.014 | 0 |  | 0.014 | 0 |
| FD&C Yellow Dye #5 | 0.04 | 0 |  | 0.04 | 0 |  | 0 | 0 |  | 0 | 0 |
| FD&C Red Dye #40 | 0 | 0 |  | 0.02 | 0 |  | 0 | 0 |  | 0.03 | 0 |
| FD&C Blue Dye #1 | 0.01 | 0 |  | 0 | 0 |  | 0.02 | 0 |  | 0 | 0 |
| *Total* | *1052.6* | *4192.0* |  | *848.8* | *4192.0* |  | *849.6* | *4192.0* |  | *854.2* | *4192.0* |

**Supplementary Table 2**: Fatty acid percent composition of the LF (Low Fat) diet containing 10% fat by weight, High Fat (HF) diet containing 45% fat by weight, and 45% HF diets supplemented with 1.6% WPPC (HF1.6%) and 10% WPPC (HF10%). Data are expressed as mean ± SD. Each diet was analyzed 4 to 5 times.

|  | LF (n=5) | | | HF (n=4) | | | HF 1.6% (n=4) | | | HF 10% (n=4) | | |
| --- | --- | --- | --- | --- | --- | --- | --- | --- | --- | --- | --- | --- |
| C10:0 | 0 | ± | 0 | 0.05 | ± | 0.005 | 0.06 | ± | 0.01 | 0.16 | ± | 0.01 |
| C12:0 | 0 | ± | 0 | 0.07 | ± | 0.004 | 0.11 | ± | 0.01 | 0.31 | ± | 0.01 |
| C14:0 | 0.8 | ± | 0.0 | 1.3 | ± | 0.1 | 1.4 | ± | 0.1 | 2.1 | ± | 0.0 |
| C16:0 | 20.7 | ± | 0.9 | 25.4 | ± | 2.0 | 25.1 | ± | 2.1 | 24.7 | ± | 0.1 |
| C16:1 | 0.8 | ± | 0.0 | 1.6 | ± | 0.1 | 1.6 | ± | 0.1 | 1.6 | ± | 0.0 |
| C18:0 | 12.9 | ± | 0.6 | 18.3 | ± | 1.5 | 18.1 | ± | 1.4 | 16.8 | ± | 0.2 |
| C18:1n9 | 26.3 | ± | 0.7 | 33.7 | ± | 2.8 | 34.4 | ± | 2.1 | 34.8 | ± | 1.0 |
| C18:1n7 | 1.7 | ± | 0.0 | 0.5 | ± | 0.9 | 0.05 | ± | 0.1 | 0.1 | ± | 0.01 |
| C18:2n6 | 33.8 | ± | 0.8 | 17.7 | ± | 1.5 | 17.8 | ± | 1.5 | 17.9 | ± | 0.7 |
| C18:3n3 | 3.0 | ± | 0.6 | 1.1 | ± | 0.2 | 1.0 | ± | 0.1 | 1.2 | ± | 0.2 |
| C20:4n6 | 0.0 | ± | 0.0 | 0.3 | ± | 0.03 | 0.2 | ± | 0.03 | 0.3 | ± | 0.03 |
| Total fatty acid concentrations (mg/g diet) | 44 | ± | 2 | 253 | ± | 61 | 258 | ± | 29 | 270 | ± | 16 |

**Supplementary Table 4**: Hippocampal Polar Lipid Fatty Acid Concentrations in Wistar Rats receiving a low fat (LF), high fat (HF), high fat + 1.6% WPPC (HF1.6) or high fat + 10% WPPC (HF10) diet. Concentrations are expressed in units of mg fatty acid per gram of brain.

| Group | **C16:0** | **C16:1** | **C17:0** | **C18:0** | **C18:1** | **18:2n6** | **18:3n3** | **20:4n6** | **20:5n3** | **22:6n3** | **Total**  **Fatty Acids** |
| --- | --- | --- | --- | --- | --- | --- | --- | --- | --- | --- | --- |
| *Sphingolipid* | | | | | | | | | | | |
| LF, n=9 | 0.028±0.006 | ND | 0.013±0.022 | 0.099±0.022 | 0.001±0.001 | ND | 0.006±0.018 | 0.018±0.005 | 0.002±0.001 | 0.005±0.003 | 0.173±0.032 |
| HF, n=10 | 0.044±0.025 | 0.003±0.004 | 0.017±0.008 | 0.125±0.053 | 0.002±0.003 | ND | ND | 0.024±0.009 | 0.007±0.006 | 0.011±0.006 | 0.232±0.096 |
| HF1.6, n=11 | 0.028±0.014 | 0.001±0.003 | 0.012±0.002 | 0.089±0.031 | ND | ND | 0.005±0.017 | 0.017±0.002 | 0.003±0.002 | 0.007±0.005 | 0.162±0.059 |
| HF10, n=8 | 0.036±0.012 | 0.007±0.015 | 0.015±0.002 | 0.105±0.034 | 0.007±0.016 | ND | ND | 0.026±0.021 | 0.012±0.027 | 0.019±0.039 | 0.230±0.150 |
| *Lyso-PC* | | | | | | | | | | | |
| LF, n=9 | 0.030±0.010 | ND | 0.11±0.002 | 0.029±0.011 | 0.004±0.001 | 0.002±0.001 | ND | ND | ND | ND | 0.075±0.017 |
| HF, n=10 | 0.037±0.022 | ND | 0.015±0.006 | 0.036±0.013 | 0.006±0.006 | 0.001±0.002 | ND | ND | ND | ND | 0.095±0.045 |
| HF1.6, n=11 | 0.024±0.005 | ND | 0.012±0.004 | 0.022±0.007 | 0.003±0.002 | 0.002±0.002 | ND | ND | ND | ND | 0.063±0.009 |
| HF10, n=8 | 0.044±0.058 | ND | 0.014±0.003 | 0.027±0.009 | 0.005±0.005 | 0.003±0.003 | ND | ND | ND | ND | 0.092±0.073 |
| *PC* | | | | | | | | | | | |
| LF, n=9 | 2.281±0.523 | ND | 0.073±0.144 | 0.676±0.134 | 1.075±0.143 | 0.071±0.113 | ND | 0.325±0.183 | ND | 0.073±0.044 | 4.772±0.888 |
| HF, n=10 | 1.978±0.823 | ND | ND | 0.660±0.158 | 1.020±0.174 | 0.060±0.101 | ND | 0.329±0.192 | ND | 0.090±0.061 | 4.343±1.341 |
| HF1.6, n=11 | 1.874±0.365 | ND | 0.052±0.172 | 0.611±0.116 | 0.913±0.147 | 0.213±0.081 | ND | 0.253±0.082 | ND | 0.065±0.040 | 4.015±0.693 |
| HF10, n=8 | 2.071±0.404 | ND | 0.003±0.008 | 0.698±0.130 | 1.004±0.215 | 0.178±0.124 | ND | 0.315±0.082 | ND | 0.072±0.047 | 4.365±0.897 |
| *PS* | | | | | | | | | | | |
| LF, n=9 | 0.057±0.051 | ND | 0.012±0.005 | 0.295±0.253 | 0.074±0.063 | 0.003±0.003 | ND | 0.043±0.038 | ND | ND | 0.491±0.328 |
| HF, n=10 | 0.076±0.083 | ND | 0.016±0.005 | 0.358±0.248 | 0.104±0.062 | 0.008±0.012 | ND | 0.070±0.062 | ND | ND | 0.640±0.338 |
| HF1.6, n=11 | 0.066±0.071 | ND | 0.064±0.175 | 0.321±0.224 | 0.104±0.070 | 0.010±0.025 | ND | 0.042±0.045 | ND | ND | 0.613±0.354 |
| HF10, n=8 | 0.046±0.013 | ND | 0.017±0.011 | 0.377±0.341 | 0.194±0.303 | 0.006±0.008 | ND | 0.042±0.019 | ND | ND | 0.711±0.444 |
| *PI* | | | | | | | | | | | |
| LF, n=9 | 0.072±0.049 | 0.013±0.006 | 0.015±0.005 | 0.340±0.192 | 0.090±0.062 | 0.006±0.003 | ND | 0.060±0.041 | ND | ND | 0.603±0.277 |
| HF, n=10 | 0.078±0.049 | 0.017±0.008 | 0.019±0.009 | 0.228±0.178 | 0.063±0.049 | 0.007±0.006 | ND | 0.060±0.036 | ND | ND | 0.477±0.250 |
| HF1.6, n=11 | 0.061±0.037 | 0.012±0.006 | 0.037±0.087 | 0.148±0.069 | 0.038±0.022 | 0.004±0.003 | ND | 0.063±0.032 | ND | ND | 0.369±0.124 |
| HF10, n=8 | 0.081±0.048 | 0.014±0.010 | 0.017±0.014 | 0.296±0.180 | 0.089±0.046 | 0.008±0.012 | ND | 0.089±0.052 | ND | ND | 0.599±0.201 |
| *PE* | | | | | | | | | | | |
| LF, n=9 | 0.243±0.093 | ND | 0.019±0.004 | 0.874±0.224 | 0.512±0.237 | 0.011±0.011 | ND | 0.562±0.273 | ND | 0.623±0.288 | 2.913±1.097 |
| HF, n=10 | 0.279±0.069 | ND | 0.024±0.006 | 0.967±0.319 | 0.611±0.171 | 0.014±0.006 | ND | 0.656±0.194 | ND | 0.732±0.235 | 3.358±0.969 |
| HF1.6, n=11 | 0.228±0.058 | ND | 0.081±0.208 | 0.754±0.262 | 0.510±0.136 | 0.012±0.005 | ND | 0.561±0.145 | ND | 0.639±0.201 | 2.841±0.772 |
| HF10, n=8 | 0.313±0.068 | ND | 0.027±0.008 | 0.940±0.226 | 0.524±0.113 | 0.025±0.024 | ND | 0.678±0.159 | ND | 0.678±0.160 | 3.248±0.650 |

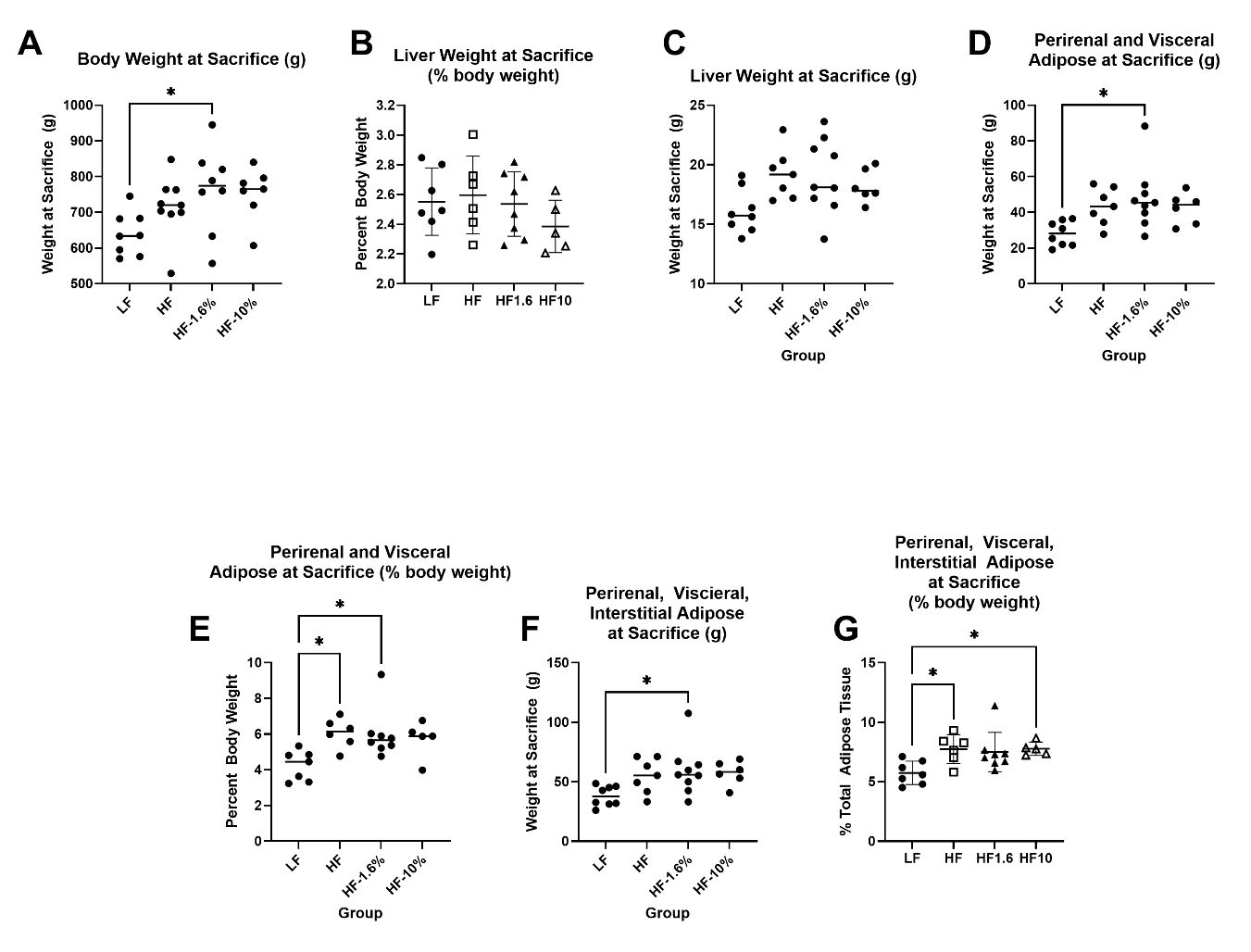

**Supplementary Figure 1**: One-way ANOVA of body weight at sacrifice (**A**), liver (**B**), perirenal and visceral adipose (**D**), and total adipose less testicular adipose (**F**) weights, and of the fraction of total body weight of liver (**C**), perirenal and visceral adipose (**E**), total adipose less testicular adipose (**G**) of male Wistar rats (LF n=9, HF n=7, HF1.6 n=10, HF10 n=6) placed on a low fat diet (LF, 10% fat by weight), high fat diet (HF, 45% fat by weight), or a high fat diet supplemented with 1.6% or 10% WPPC by weight (HF1.6% and HF10% respectively).

**Supplementary Figure 2**: Representative example measurement of an evoked response in a Wistar Rat on the high fat diet supplemented with 1.6% whey protein phospholipid concentrate. The stimulus is highlighted in blue (**A**), the field potential is highlighted in green (**B**), and the evoked response is highlighted in purple (**C**).

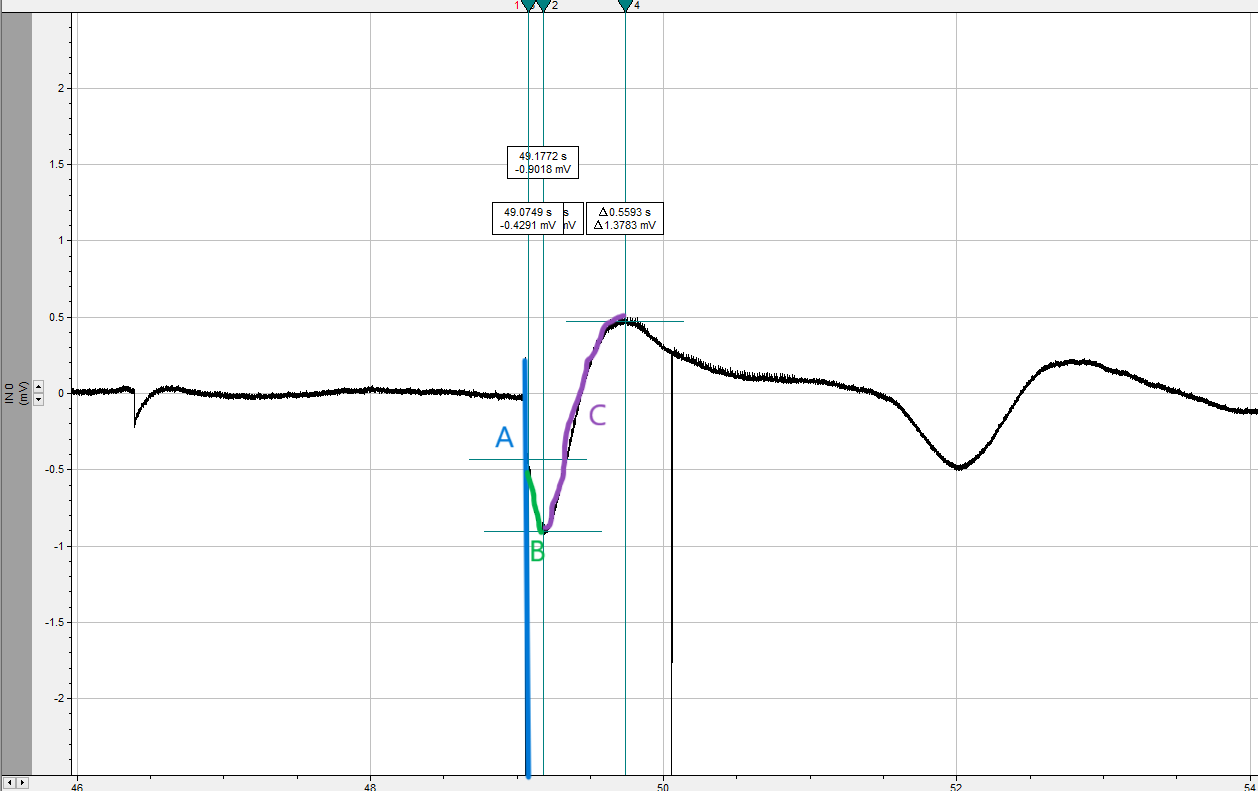

**Supplementary Figure 3**: Comparison of the 50% evoked response (ED50) (**A**) and the ratio of evoked response at 30-60 minutes to baseline (**B**) in rats consuming a LF, HF, HF1.6 or HF10 diet. There were no significant differences in the ED50 or the ratio by one-way ANOVA.

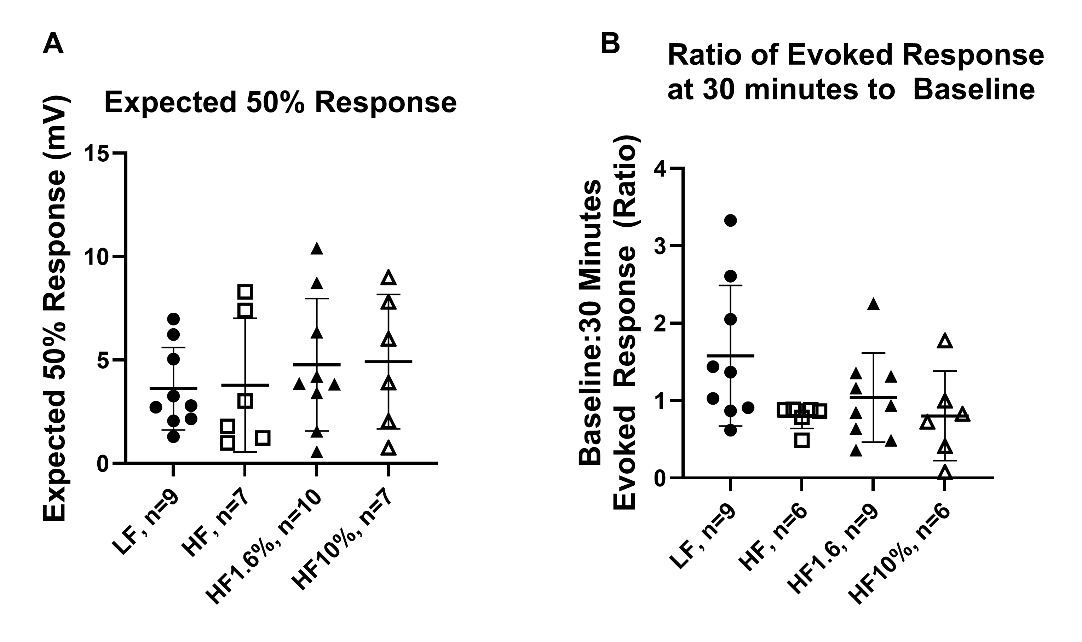
